# Probing inhomogeneous diffusion in the microenvironments of phase-separated polymers under confinement

**DOI:** 10.1101/402230

**Authors:** Marjan Shayegan, Radin Tahvildari, Lydia Kisley, Kimberly Metera, Stephen W. Michnick, Sabrina R. Leslie

## Abstract

Biopolymer condensates formed by liquid-liquid phase separation of proteins and nucleic acids have been recently discovered to be prevalent in biology. These dynamic condensates behave like biochemical reaction vessels but little is known about their structural organization and properties. Their biophysical properties and catalytic functions are likely related to condensate size, and thus it is critical that we study them on scales found *in vivo*. However, previous *in vitro* studies of condensate assembly and physical properties have involved condensates up to 1000 times larger than those found *in vivo*. Here, we report the application of confinement microscopy to visualize condensates and control their sizes by creating appropriate confinement length scales relevant to the cell environment. We observe anomalous diffusion of probe particles embedded within confined condensates, as well as heterogeneous dynamics in condensates formed from PEG/dextran and in ribonucleoprotein complexes of RNA and the RNA-binding protein Dhh1. We propose that the non-Gaussian dynamics we observe may indicate a *hopping diffusion* mechanism inside condensates. We also observe that for dextran-rich condensates, but not for ribonucleo condensates, probe particle diffusion depends on condensate size.

---

Liquid-liquid phase separation is found throughout nature and has important consequences in biology (1–3). For example, biopolymer condensates, or non-membranous organelles (NMOs), are dynamic structures within the cyto- and nucleoplasm of cells that form *via* reversible and highly controlled liquid-liquid phase separation of proteins and nucleic acids. Inside cells, different NMOs perform a variety of functions (3–8): concentrating molecules to enhance biochemical reaction rates within the diverse and crowded cell interior; isolating catalytic processes; sequestering and storing materials for later release; and helping cells respond to stress. Some are further sub-compartmentalized and may generate gradients and chemical potentials (5). Some appear to concentrate specific molecules while simultaneously excluding others. Through aberrant formation, regulation, or stability, NMOs may also contribute to neurodegenerative conditions including Alzheimer’s and Huntington’s diseases. However, the principles governing NMO assembly, disassembly and functions remain unknown (9–12). Furthermore, the relationships between NMO size, properties, and functions need to be clarified. NMOs scale with the size of the cell (4, 13), and cell size dysregulation is connected to many diseases, including cancers (14).

Detailed biophysical studies of phase-separated biopolymer condensates will help clarify how these important structures function. Important factors for the phase separation of biopolymer condensates include: 1) the content of intrinsically disordered regions in the component proteins, and 2) interactions between folded protein or RNA domains with linear regions (15). The mechanical properties of the phase separated condensates (e.g. surface tension and viscoelastic properties) and the interactions among their component proteins and nucleic acids are expected to control condensate coalescence, internal structuring, diffusion within the condensate, and exchange of molecules with the cytoor nucleoplasm, and thus must heavily influence condensate function (3). For example, it has been proposed that the viscoelastic properties of NMOs control enzymatic processes by lowering the flux of specific molecular species (16).

*In vitro* studies of biopolymer condensates allow for controlled conditions in which to study condensate assembly and properties, and have probed a range of parameters controlling phase separation (Refs.(3, 17–20)are just a few examples of recent reviews describing insights from *in vitro* studies). However, typical *in vitro* techniques cannot approach simulating the naturally crowded, complex, and confined conditions found in cells, and reconstituted condensates can vary greatly in size and be up to 10-1000 times larger (*∼*10 *-*100 *μ*m) than those observed *in vivo* (*∼*100 nm for P-bodies (7)). To understand natural NMOs, it is crucial to determine how these vastly different size regimes might influence condensate biophysical properties.

## Experimental approach

Convex Lens-induced Confinement (CLiC) fluorescence microscopy (21–24) is applied to control condensate size and monitor how size affects diffusion within them. In this technique, condensates are formed within tiny wells (450 nm deep and 3-50 *μ*m in diameter) to limit condensate growth. In a typical confinement microscopy experiment, a fluid sample of biomolecules is loaded into a custom-designed imaging chamber made from two glass coverslips separated by a spacer. As shown in Fig. 1, the bottom coverslip (the chamber floor) contains an array of embedded wells with highly-controlled and well-defined nano-to microscale geometry. After sample injection, the deformable glass roof is pressed downwards by a convex lens, gently herding the biomolecules into the wells. The floor-ceiling distance can be controlled by pillars/posts attached to the chamber ceiling to leave a small gap by which to introduce molecules/reagents to the large, trapped biomolecules. The wells are deep enough to contain freely diffusing and fluctuating biomolecules, but shallow enough to keep them in the field of view for extended observation times. By squeezing out extra molecules, background fluorescence is reduced up to 20-fold and samples can be tested at higher concentrations compared to typical single-molecule techniques. Furthermore, the array of embedded wells allows data collection with significant statistics within a single experiment.

**Fig. 1.**
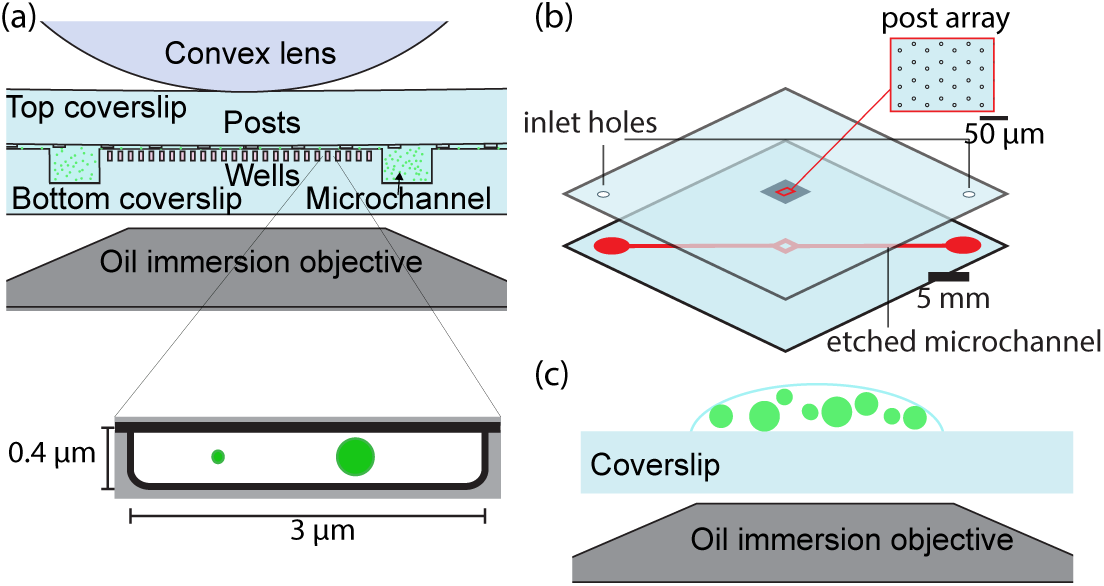
Confinement microscopy: (a) schematic of a Convex Lens-induced Confinement (CLiC) instrument used here; a convex lens deforms the top coverslip of the flow cell. Posts fabricated into the top coverslip maintain a space above the wells (450 nm deep and 3 *μ*m in diameter) embedded in the flow cell floor, where condensates are formed).(b) Schematic of a microchannel fabricated into the bottom coverslip that encircles the embedded pit array and facilitates reagent addition and buffer exchange. (c) Schematic showing condensate formation on an open surface, i.e. no confinement.

Confinement microscopy enables a new approach to explore macromolecular behavior, specifically protein phase separation, on various dimensions under spatial confinement relevant to the cellular interior. We visualized formation of condensates and controlled their growth by systematically varying confinement dimensions (Fig. 2 and Fig. 3). We then explored the mesoscale properties of confined condensates by tracking the fluctuating motions of encapsulated fluorescent probe particles within the condensates. We investigated two liquid condensate systems. First, phase-separated polyethylene glycol (PEG) and dextran condensates were used to explore assembly, growth and final size under confinement. PEG-dextran condensates have been previously formed inside large vesicles, a conventional technique, as a simple model of the crowded cytoplasm (25–28). Second, we investigated biopolymer condensates of ribonucleoprotein complexes formed of polyuridine (polyU) RNA and protein Dhh1 as a simple, few-component model of natural NMOs. In both cases, we explored probe particle diffusion with respect to condensate size.

**Fig. 2.**
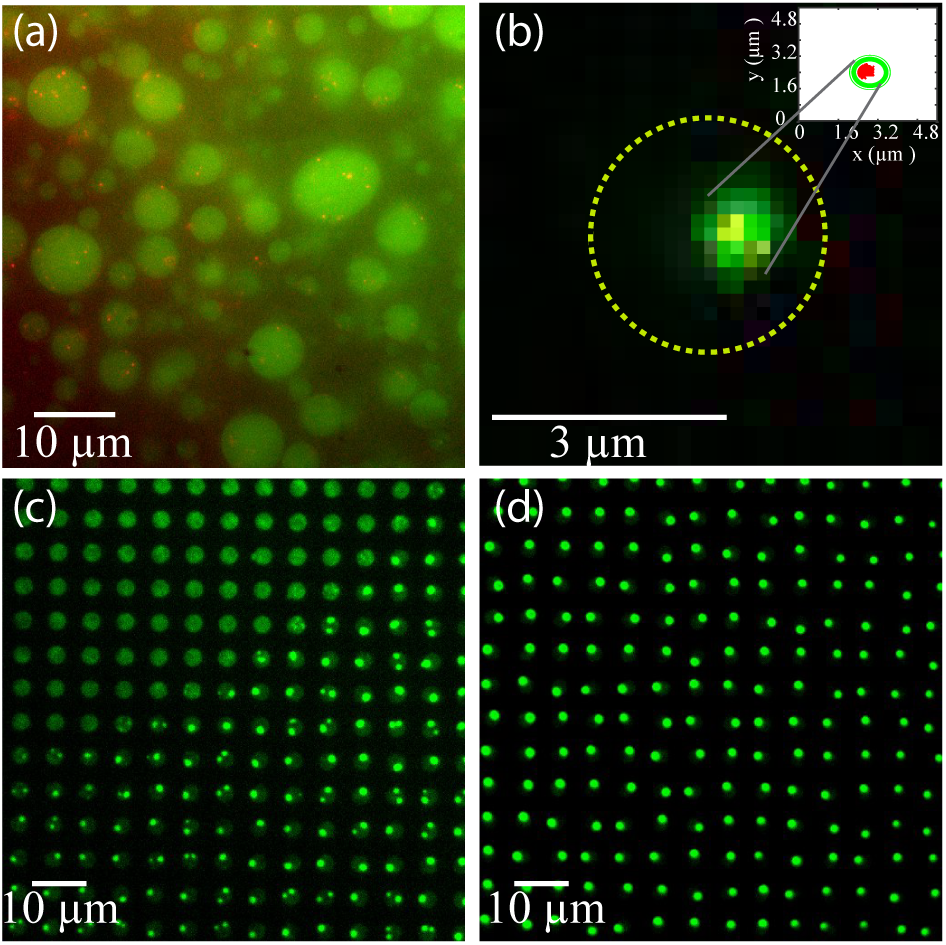
Dextran-rich condensates condensed from a mixture of PEG-dextran formed (a) on an open surface (i.e. without confinement), and (b) inside a 3 *μ*m well (i.e. under confinement). Using dual color imaging, fluorescently-labeled beads (red) are distinguished in dextran-rich condensates (green). Note: Red beads appear yellow in the green channel when they overlap with green-labeled dextran-rich droplets. Dashed circle: well location. Inset: the particle tracking of a red-labeled bead inside the condensate (green circle). (c) Initiation of phase separation in an array of wells containing confined labeled dextran, as PEG is introduced from the bottom right corner. (d) Same array as shown in (c) 5 minutes later, condensates reach their final size.

**Fig. 3.**
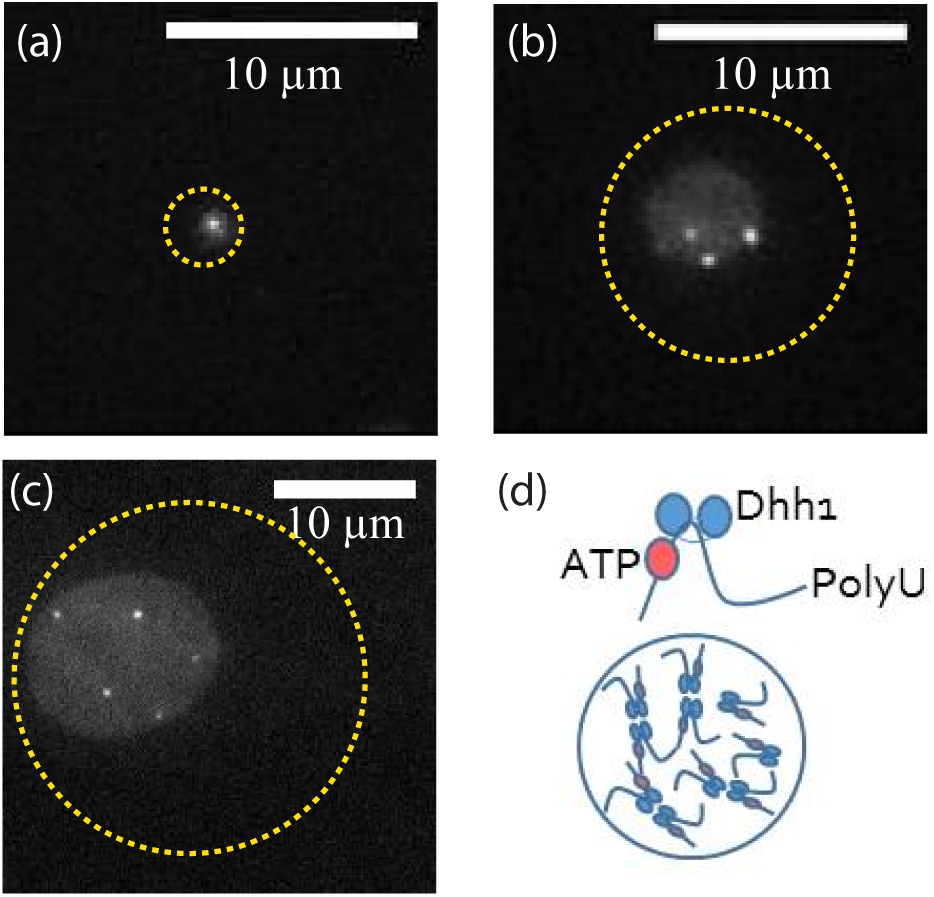
Effect of confinement on the final size of the protein condensates: condensate of (a) 2 *μ*m, (b) 4 *μ*m, and (c) 14 *μ*m formed in wells with diameters of 3, 10, and 25 *μ*m, respectively. Dashed circles correspond to the location of the wells. Condensates are shown in gray and bright dots inside condensates are probing particles of 48 nm in diameter. (d) Schematic of ATP- and RNA-bound Dhh1 conformation and how these components may interact with each other inside a phase separated condensate.

## PEG-dextran mixtures exhibit anomalous diffusion and size-dependent properties

In the first system, fluorescently-labeled dextran was confined within wells, and condensate formation and growth were monitored as PEG was introduced. The condensates formed under confinement (see Supplementary info S1 for Materials and Experimental Methods) were demonstrably smaller and more monodisperse than those formed on a featureless coverslip where condensate fusion may occur more quickly (Fig. 1c and Fig. 2a). When the reconstituent elements are confined in sealed wells, the maximum size of the phase separated condensate is controlled and limited by the amount of material available inside the well, *i.e.* proportional to the well size. For example, Fig. 2 compares condensates formed on a coverslip and within a large array of wells with a controlled final size of the condensates.

Previous work indicated that smaller condensates may have different network structure and physical behaviors compared to larger ones such that some mechanical theories would fail to explain their observed behavior (29, 30). To test whether the properties of confined condensates change with their size, we performed passive microrheology experiments. The motions of probe particles (20-nm fluorescent latex microspheres passivated with dextran), randomly distributed within condensates, were tracked using our in-house tracking algorithm (31) (Fig. 2b; Supplementary Info S2). To characterize diffusion properties of the embedded probe particles, we calculated their time-averaged mean squared displacements (MSD) (5), and performed correlation analyses on the videos of particle diffusion. Correlation methods such as fluorescence correlation spectroscopy (32, 33) and image correlation spectroscopy (34) are complimentary to single particle tracking, but are more objective in that individual point spread functions do not need to be fit. As such, correlation techniques are able to quantify faster diffusion coefficients or assess data with lower signal to noise ratios (35). The self-similarity of the fluorescent intensity signal at the pixel with the highest intensity fluctuations within the condensate was analyzed by second-order correlation (see Supplementary Info S3 for more details).

For solutions of pure dextran prior to PEG introduction, a homogeneous diffusion of beads was obtained in the microscale (Supplementary Figure S1). Interestingly, after PEG was introduced and phase separation occurred, dextran-rich condensates possessed heterogeneous properties (Fig. 4(a)). Probe particle diffusion varied from one condensate to another, and even from one location to another within the same condensate; MSD values varied by up to one order of magnitude.

**Fig. 4.**
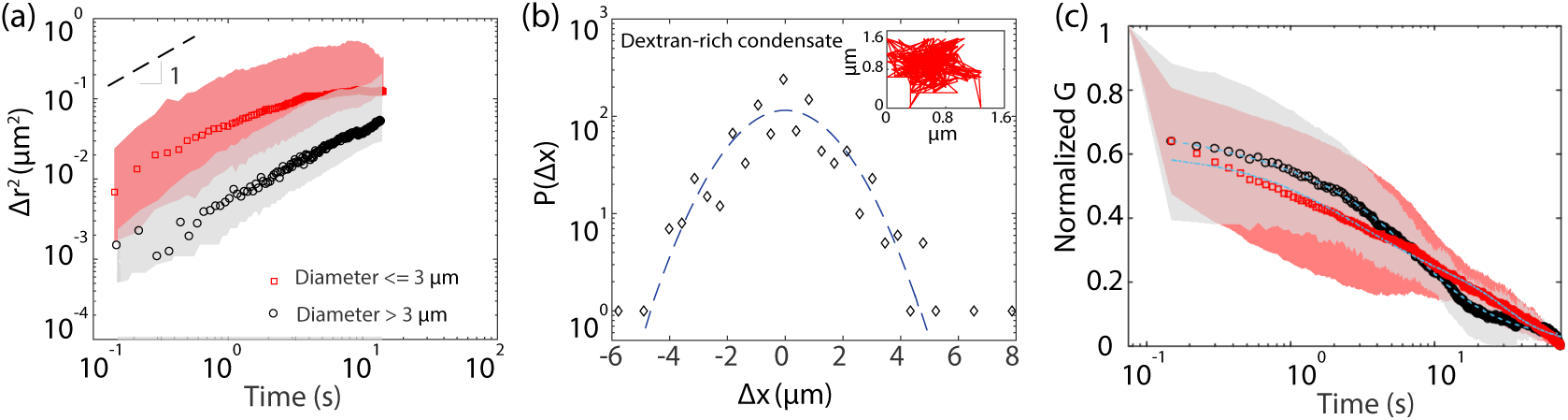
Diffusion analysis for probe particles inside PEG-dextran condensates (N=135): Red and black represent condensates that are smaller and larger than 3 *μ*m, respectively. (a) Time-averaged mean squared displacement versus lag time of probe particles embedded in dextran-rich condensates. For each data set, the circles or squares show the averaged MSD (i.e. time and ensemble averaged), and the shaded area the distribution of MSD curves within that set. Dashed line: a slope of 1 in log-log scale, indicative of normal Brownian diffusion. (b) Typical displacement probability distribution, measured for a probe particle embedded inside a dextran-rich condensate, plotted logarithmically against displacement. The dashed curve represents a fitted standard Gaussian distribution, and highlights the poor fit to Brownian solutions with the experimental data. Inset: probe particle trajectory over 1 minute. (c) Correlation analysis performed on probe particle diffusion inside a dextran-rich condensates. Circles or squares show the averaged correlation curve, G, for each data set, while the shaded areas show the distribution of correlation curves. Lines represent the fit which give two decay times (relevant to a slow and a fast diffusion mode) and power law exponents.

To help elucidate the diffusion mechanisms within the PEG-dextran condensates, we consider the concentration regimes of the system. Table 1 shows the theoretically calculated overlap concentration (*C*^*\**^), above which polymer chains start to interact and become an entangled network, and the estimation of the dextran concentration within a condensate (C_condensate_). Overlap concentration is calculated based on 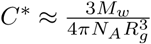, where *M*_*w*_ is molecular weight of the polymer (500 kDa for dextran and 57.5 kDa for Dhh1), *N*_*A*_ is Avogadro’s number, and *R*_*g*_ is radius of gyration of the polymer (*∼* 20*nm* for dextran and *∼* 4.8*nm* for Dhh1). Comparison suggests that a dextran-rich condensate is indeed a crowded, highly entangled system (C_condensate_>C*). Crowded environments created by long chains of dextran molecules (500 kDa) within a condensate may result in internal structural heterogeneity, leading to the observed heterogeneous diffusion behavior.

**Table 1.** Comparison of overlap concentration with polymer concentration within a condensate for a dextran-rich condensate and Dhh1-rich condensate

| Concentration | C* | C<sub>initial</sub> | C<sub>condensate</sub> | Comparison |
| --- | --- | --- | --- | --- |
| Dextran (wt%) | 1.8 | 2 | >2 | C<sub>condensate</sub>>C* |
| Dhh1 (mM) | 3.5 | 0.012 | 0.125 | C<sub>condensate</sub><C* |

Particle motion is also affected by the confining surface (*i.e.* condensate-solvent interface) through solvent-mediated hydrodynamic interactions (36). As a particle moves inside a condensate, it propagates perturbations in the velocity field of the medium traveling back and forth between the interface and the particle. In a crowded environment, hydrodynamic interactions become even more complex and can create heterogeneity in particle mobility within a condensate.

PEG-dextran condensate properties appear to be size-dependent. Despite heterogeneity, ensemble averaged MSD curves show reduced mobility of probe particles inside smaller (1-3 *μ*m) condensates compared to inside larger (>3-15 *μ*m) ones (Fig. 4 (a)). Correlation analysis (Supplementary Info S3) further supports the size dependence and heterogeneous diffusive motion of beads inside dextran condensates (Fig. 4(c)). According to the correlation analysis, particles have slower diffusion in the smaller condensates (see Table S1), similar to the results obtained from single particle tracking analysis.

## Ribonucleo poly(U)-Dhh1 condensates also exhibit anomalous diffusion

The next condensates we studied were the simple model NMOs formed from the yeast RNA binding protein Dhh1 and the RNA analogue poly(U). Here, we premixed the Dhh1 with the poly(U), and visualized formation of Dhh1-poly(U) condensates with controlled sizes using confinement microscopy (Fig. 3).

As in the PEG-dextran system, both particle tracking and correlation analysis (Fig. 5) indicated heterogeneous anomalous diffusion of embedded probe particles (48-nm fluorescent latex microspheres passivated with bovine serum albumin, BSA). However, unlike for the PEG-dextran condensates, the Dhh1-poly(U) condensates did not exhibit significant size-dependent internal properties. In addition, the averaged correlation curves were better fit with a two-component correlation function, *i.e.* two diffusion components (Fig. 5c and Supplementary Information S3), suggesting both a slow and a fast mode of diffusion. The faster component had an anomalous factor, *α*, >1 indicative of a super-diffusive, hopping mechanism.

**Fig. 5.**
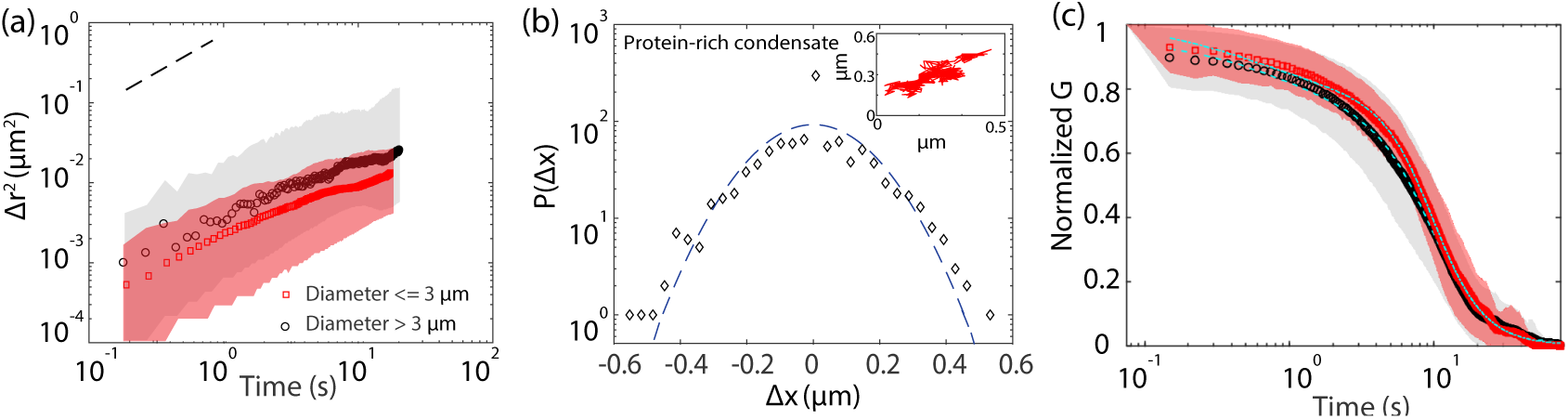
Diffusion analysis for probe particles inside protein-nucleic acid condensates (N=140): Red and black represent condensates that are smaller and larger than 3 *μ*m, respectively. (a) Time-averaged mean squared displacement versus lag time of probe particles embedded in protein-rich condensates. (b) Typical displacement probability distribution, measured for a probe particle embedded inside a protein-rich condensate plotted logarithmically against displacement. The non-Gaussianity observed can be attributed to the hopping diffusion. Inset: the trajectory of a probe particle over 1 minute. It suggests that the particle spends most of its time in a cage formed by its neighboring chain structures while during rare events moves a significant distance due to a rearrangement of cage structure. (c) Correlation analysis performed on probe particles diffusing inside a protein-rich condensate.

Anomalous diffusion and hopping mechanisms have recently been reported *in vivo* in stress granules, a different type of NMO (37). In that case, biphasic partitioning of biopolymers resulted in their having suppressed diffusion in some local microenvironments but larger mobility at other locations (38).

Single-component Brownian curves failed to fit the data (Supplementary Figure S2). Similar to the single-particle tracking results, there were statistically similar values for the Dhh1-poly(U) condensates regardless of size (Supplementary Table S1).

In order to better understand the anomalous (non-Brownian) diffusion within condensates, we measured the displacement probability distribution. Compared to a Gaussian model, there are too many large displacement events (the tails in Fig. 5(b) at high and low Δ*x* values) and a stronger confinement of motion (the sharp peak close to zero displacement, Δ*x* = 0). These observations provide further evidence for the “hopping diffusion” mechanism where a particle mostly resides in a caged structure made by neighboring molecules but occasionally escapes and moves a significantly larger distance than a normal Brownian diffusion would predict (38, 39). The non-Gaussian distribution of particle displacement was also observed in dextran condensates (Fig. 4(b)) with tails at long displacements and is an indication of a heterogeneous environment, although unlike in protein condensates, hopping diffusion is not directly observed from analysis of particle trajectory (inset in Fig. 4b).

To explain the diffusion mechanisms within the protein condensates, protein concentrations inside condensates were estimated. The estimated Dhh1 concentration (see Supplementary Figure S3) inside a condensate and the theoretical overlap concentration for the protein condensates are shown in Table 1. Comparison of these values suggests that Dhh1 protein itself does not create a significantly crowded condensate interior, yet we still observe confined motion of embedded probe particles. Poly(U) molecules may be unevenly distributed within the condensate to cause locally crowded environments that inhibit particle motion. It is also possible that specific interactions between the condensate components contribute to the condensate interior properties and effectively suppress particle movement. Binding of ATP and RNA to Dhh1 has been previously shown to be necessary for condensate formation, presumably due to inducing specific conformations of the Dhh1 protein (6). A schematic of this bound conformation of poly(U) and Dhh1 is shown in Fig. 3(d).

To explore the effect of Dhh1-/poly(U) bound conformation in condensate assembly, we introduced the regulatory protein Not1 to the confined condensates. Not1 enhances the ATPase activity of Dhh1, unbinds Dhh1 from the nucleic acid by changing the Dhh1 conformation, and ultimately results in disassembly of the condensate (6). Interestingly, we observed that when we added Not1, embedded beads started to diffuse more quickly and eventually diffused freely throughout the pit in the solution of the poly(U) and Dhh1 (Supplementary Figure S4). Our particle tracking measurements showed that in an ATP- and RNA-bound Dhh1 conformation, i.e. assembled condensate, probe diffusion was highly suppressed and non-homogeneous across the condensate, but as the Dhh1 conformation changed due to the presence of Not1, probe particles diffused more freely and uniformly.

Other results have indicated that permeability and diffusion inside a protein condensate are affected by the length (MW) of the component RNA (40). These specific effects will be dictated by the type of RNA-protein interactions. Short RNAs may modulate protein-protein interactions to lead to a lowering of the viscosity, while larger molecules have the opposite effect (41), perhaps because larger molecules are involved in additional physical processes such as entanglement. In our protein study, the model RNA molecule, poly(U), is polydisperse in size (100kDa < MW < 1000 kDa), making it potentially more biologically representative than a monodisperse RNA molecule. This polydispersity likely enhances the heterogeneity of the condensate interior structure, as indicated by our observed particle diffusion.

In summary, this study showcases a new application of confinement microscopy, which has enabled 1) control of condensate size through spatial confinement (i.e. well with a particular size), as shown in Fig. 3(a-c), and 2) the quantitative characterization of micro-environments within condensates. We obtained heterogeneous dynamics as well as anomalous diffusion inside both dextran-rich and protein-rich condensates. In ribonucleo protein-rich condensates, we found evidence of temporal heterogeneity such that embedded particles may diffuse within a microenvironment and occasionally “hop” to a neighboring microenvironment with different physical properties. Furthermore, we observed a size dependence of diffusion, and thus interior condensate properties, for dextran condensates, although this was not observed significantly for the Dhh1-poly(U) condensates tested. The presence of microenvironments inside NMO condensates has important biological implications. Proposed to arise in part due to heterogeneous distribution of nucleic acids within a condensate, microenvironments likely facilitate biochemical processes such as gene expression (42). Molecular crowding also contributes to the stochasticity of biochemical processes within NMOs by decreasing diffusion of proteins and nucleic acids and contributing to the uneven distribution of biomolecules (42). *In vitro* studies at *in vivo* size scales should shed light on important condensate properties. Confinement microscopy is a promising technique for a variety of NMO investigations, providing insight into single molecule dynamics and reaction kinetics, by creating “cell-like” conditions of spatial confinement and crowding.

## Acknowledgements

M.S., S.W.M, and S.R.L. acknowledge support from the Canadian Institutes of Health Research (CIHR) grants MOP-GMX-152556 (S.W.M.), the Human Frontier Science Program RGP0034/2017 (S.W.M.), the Natural Sciences and Engineering Research Council grant, Discovery and Accelerator Grant Programs (S.R.L.), and CREATE Program on the Cellular Dynamics of Macromolecular Complexes (M.S.). L.K. thanks the Arnold O. and Mabel M. Beckman Foundation for support by the Beckman-Brown Interdisciplinary Postdoctoral Fellowship. The authors thank Karsten Weis and Marie Hondele (ETH-Zurich) for gifts of Dhh1 and Not1 proteins and also thank Christine Desroches for supporting protein condensation experimental design.

